# A postmeiotic route to polyploidy

**DOI:** 10.64898/2026.02.20.706944

**Authors:** Cintia Gómez-Muñoz, Nina Vittorelli, Maxime Gaudin, Nicolas Agier, Stéphane Delmas, Yann Corbeau, Marco Cosentino Lagomarsino, Gianni Liti, Bertrand Llorente, Gilles Fischer

## Abstract

Polyploidy, the presence of multiple chromosome sets within a genome, is widespread and has played a major role in genome evolution in plants, fungi, and animals. Although polyploidization is generally attributed to mitotic genome doubling or unreduced gamete fusion, the mechanisms underlying ploidy increases, particularly triploid formation, remain unclear. Here, we identify a novel route to autopolyploidization in *Saccharomyces cerevisiae*, termed Sporulate– Endoreplicate–Mate (*SEM*), in which one gamete, or spore, undergoes postmeiotic endoreplication and subsequently mates with a nearby non-endoreplicated sibling spore after ascus breakdown, increasing ploidy by one haploid genome. We experimentally demonstrate stepwise transitions from diploidy to triploidy and from triploidy to tetraploidy. Intact asci can spontaneously generate new triploid and tetraploid strains upon one or two successive *SEM* cycles. Natural polyploids exhibit genomic signatures consistent with *SEM* being a major natural route to polyploidization in yeast. *SEM* establishes triploids as central intermediates in yeast polyploidization, analogous to the triploid bridge described in plants, and raises the possibility that a similar mechanism may contribute to polyploid genome formation in other eukaryotes.

## Introduction

Polyploidy, defined as the presence of more than two complete sets of homologous chromosomes, is widespread across eukaryotes, with both ancient and recent polyploidization events documented in all major lineages (Morris *et al*, 2024). In plants and fungi, polyploidy promotes genomic and phenotypic diversification, facilitates adaptation to stress, and contributes to ecological success and domestication (Selmecki *et al*, 2015; Albertin & Marullo, 2012; Segraves & Anneberg, 2016; Wood *et al*, 2009). Although less common in animals, polyploidy has also played an important evolutionary role, notably through two ancient rounds of whole-genome duplication (WGD) at the base of vertebrate evolution (MABLE, 2004; Ohno, 1970), and remains prevalent in fish, amphibians, and some invertebrates (Mable *et al*, 2011; David, 2022; Mezzasalma *et al*, 2023). Polyploidy can also arise within specific tissues of multicellular organisms (Morris *et al*, 2024) and is a hallmark of cancer, where genome doubling promotes genome instability genome instability, tumor evolution, and therapy resistance (Bielski *et al*, 2018; Lambuta *et al*, 2023; Martínez-Jiménez *et al*, 2023). In vertebrates, genes retained from ancient WGD (ohnologs) are also disproportionately enriched among genes implicated in cancer and dominant genetic disorders (Singh *et al*, 2012). Overall, polyploidy is a major source of genetic innovation and is increasingly recognized as a multifaceted force shaping evolution, adaptation, and disease (Morris *et al*, 2024; Otto & Whitton, 2000; Ohno, 1970).

Autopolyploidization, in which additional chromosome sets arise within a species, can occur through diverse mitotic or meiotic mechanisms (Morris *et al*, 2024; Otto & Whitton, 2000; Ramsey & Schemske, 1998). Mitotic mechanisms that result in WGD duplicate the entire genome of a cell and therefore generate even ploidy levels, such as the transition from diploidy to tetraploidy. Meiotic restitution is the omission or failure to complete one of the two meiotic divisions, producing unreduced gametes that retain the parental chromosome number rather than the normal gametic complement. First-division restitution (FDR) results from failure of meiosis I, whereas second-division restitution (SDR) results from omission or failure of meiosis II. Fusion of unreduced gametes with reduced or other unreduced gametes can generate offspring with different ploidy levels. Although triploidy has often been viewed as an evolutionary dead end because of reduced fertility, studies in plants have shown that triploids can serve as a bridge in the transition from diploidy to tetraploidy (Ramsey & Schemske, 1998; Otto & Whitton, 2000). Whether such stepwise transitions contribute to polyploid evolution remains unclear.

The budding yeast *Saccharomyces cerevisiae* provides a striking example of this problem. In the 1,011-genome study, 91 of the 742 isolates with assigned natural ploidy were polyploid (11.5%), including 46 triploids and 39 tetraploids, indicating that triploids were slightly more common than tetraploids (Peter *et al*, 2018). This abundance of triploids is difficult to reconcile with the canonical yeast life-cycle (Fischer *et al*, 2021). Diploid cells carry the two mating types, *MATa* and *MAT*_α,_ and normally undergo meiosis and sporulation to produce four haploid spores, two *MATa* and two *MAT*_α,_ which remain enclosed within an ascus. Upon germination, spores of opposite mating types can mate with one another, thereby restoring the diploid state. Despite early reports of triploid and tetraploid isolates more than seventy years ago (Pomper *et al*, 1954), and laboratory demonstrations that triploids can be generated through mating-based routes (Gunge & Nakatomi, 1972), how triploids arise in natural populations remains unknown.

A prevailing model, defined here as saltational, posits that *S. cerevisiae* undergoes or one-step transitions from diploidy to tetraploidy through two possible routes: WGD during vegetative growth or meiotic restitution followed by mating between two unreduced spores originating from the same ascus (Fig. 1A). The first route is supported by observations of spontaneous autotetraploidization during vegetative growth (Marsit *et al*, 2021; Charron *et al*, 2019; Tong *et al*, 2025). Studies of laboratory-generated polyploids have reported that newly formed tetraploids can be unstable and undergo ploidy reduction through chromosome loss, sometimes giving rise to triploid or near-triploid intermediates (31–33). In contrast, another study found that laboratory-generated tetraploids remained stable during long-term propagation, arguing against the idea that triploids are transient intermediates during ploidy regression (Lu *et al*, 2016). Importantly, ploidy stability is not restricted to laboratory generated polyploids: long term vegetative propagation of “natural polyploids,” a category that here includes both wild isolates recovered from wild niches and domesticated isolates originating from industrial or other anthropogenic environments, also revealed no reduction in ploidy, indicating that stable polyploid states can persist among natural polyploids (Dutta *et al*, 2022).

**Figure 1:**
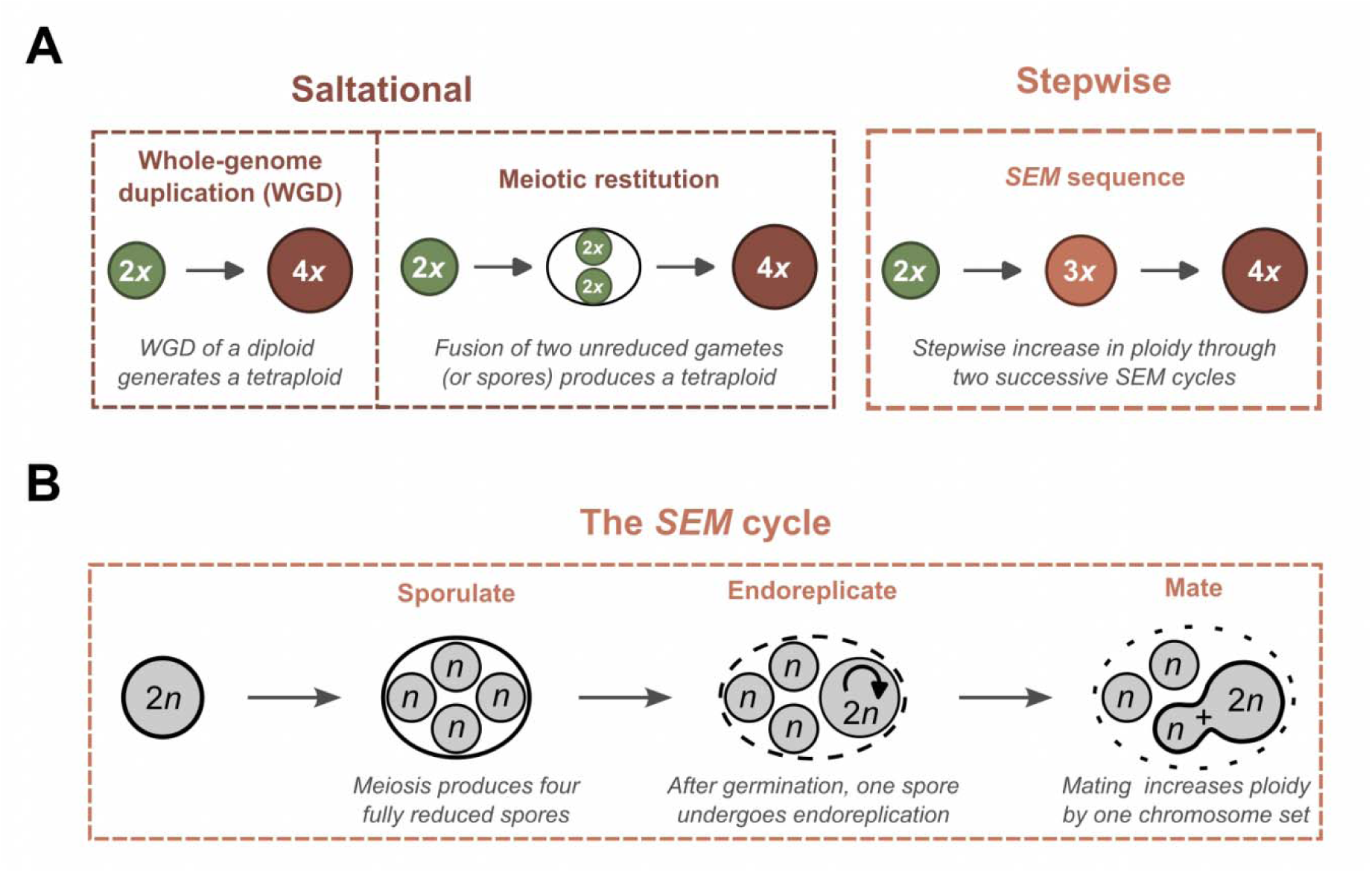
**Saltational and stepwise routes to polyploidy in *S. cerevisiae***. **A**, Schematic comparison between saltational and stepwise polyploidization, where *x* denotes ploidy level. In the saltational model, diploid cells (2*x*) transition directly to tetraploids (4*x*) in a single step, either through mitotic whole-genome duplication (WGD; left) or through meiotic restitution producing unreduced (2*x*) spores that generate tetraploids upon mating. In the stepwise polyploidization model, ploidy increases by one chromosome set at a time through successive Sporulate–Endoreplicate–Mate (*SEM*) cycles. The saltational route predicts few triploids, whereas the stepwise route predicts that triploids should arise frequently as intermediate states. **B**, Schematic representation of the *SEM* cycle, where *n* denotes the chromosome number of a gamete and 2*n* that of the corresponding vegetative cell; these terms do not indicate ploidy. Sporulation of a 2*n* vegetative cell produces four 1*n* spores enclosed within an ascus. Upon germination, one spore undergoes postmeiotic genome doubling (endoreplication) while retaining mating competence. The resulting 2*n* spore then mates with a compatible, non-doubled 1*n* spore from the same ascus, incrementing ploidy by one chromosome set.

An alternative model is stepwise polyploidization through successive gains of one chromosome set. In this model, triploids arise directly from diploids through mating with haploids, rather than as derivatives of tetraploids (Fig. 1A). This model is consistent with the relatively high frequency of triploids observed among natural polyploids (Peter *et al*, 2018; Loegler *et al*, 2024). However, such a route may appear unlikely because ordinary diploid cells do not mate, whereas haploid cells exist only transiently between spore germination and mating.

Here, we experimentally demonstrate stepwise transitions from diploidy to triploidy and from triploidy to tetraploidy in *S. cerevisiae*. We further show that natural polyploids display genomic features consistent with stepwise polyploidization. Together, our findings identify a previously unrecognized route to polyploidization that may contribute substantially to the emergence of polyploid yeast populations and may also operate in other eukaryotes.

## Results

### The Sporulate-Endoreplicate-Mate sequence

In this study, we propose a mechanism for stepwise increases in ploidy. The *SEM* cycle consists of three successive events (Fig 1B). It begins with *S,* sporulation of a 2*n* vegetative cell which produces four 1*n* gametes in the form of spores that remain together within an ascus. This is followed by *E*, a genome-doubling event (endoreplication) that occurs after meiosis, during spore germination (Fox & Duronio, 2013; Edgar & Orr-Weaver, 2001). The molecular mechanism underlying this genome doubling event is currently unknown. Here, we use *endoreplication* as a general term for genome doubling that may result either from two successive rounds of DNA replication without mitosis or from an abortive mitosis or failed cytokinesis (Fox & Duronio, 2013). Importantly, the the resulting endoreplicated 2*n* spore retains a single mating type and therefore remains capable of mating, while another 1*n* spore produced by the same meiotic event can provide a nearby, compatible mating partner. The final event is *M,* mating between the endoreplicated 2*n* spore and a compatible 1*n* spore from the same ascus, producing a cell with one additional chromosome set (+1*x*) (Fig. 1B). Each iteration of the *SEM* cycle increases ploidy by one level (Fig. 1A).

### First *SEM* cycle: stepwise triploidization

The first iteration of the *SEM* cycle drives the diploid-to-triploid transition. It starts with *S1,* sporulation of a diploid parental cell and is directly followed by *E1* the endoreplication of one of the germinating spores (Fig. 2A). Previous studies showed that mitotic endoreplication occurs frequently during haploid vegetative growth, with approximately 17% of cells spontaneously becoming diploid after 100 generations in the absence of external stress, and this frequency can reach 100% in the presence of ethanol or potassium chloride (Harari *et al*, 2018). Diploidization has also been observed during experimental evolution under glucose limitation, suggesting that changes in ploidy can arise during adaptation to nutrient-limited environments (Venkataram *et al*, 2016). We therefore asked whether endoreplication can also occur after meiosis, during spore germination, as proposed for the *E1* step of the *SEM* sequence (Fig. 2A).

**Figure 2:**
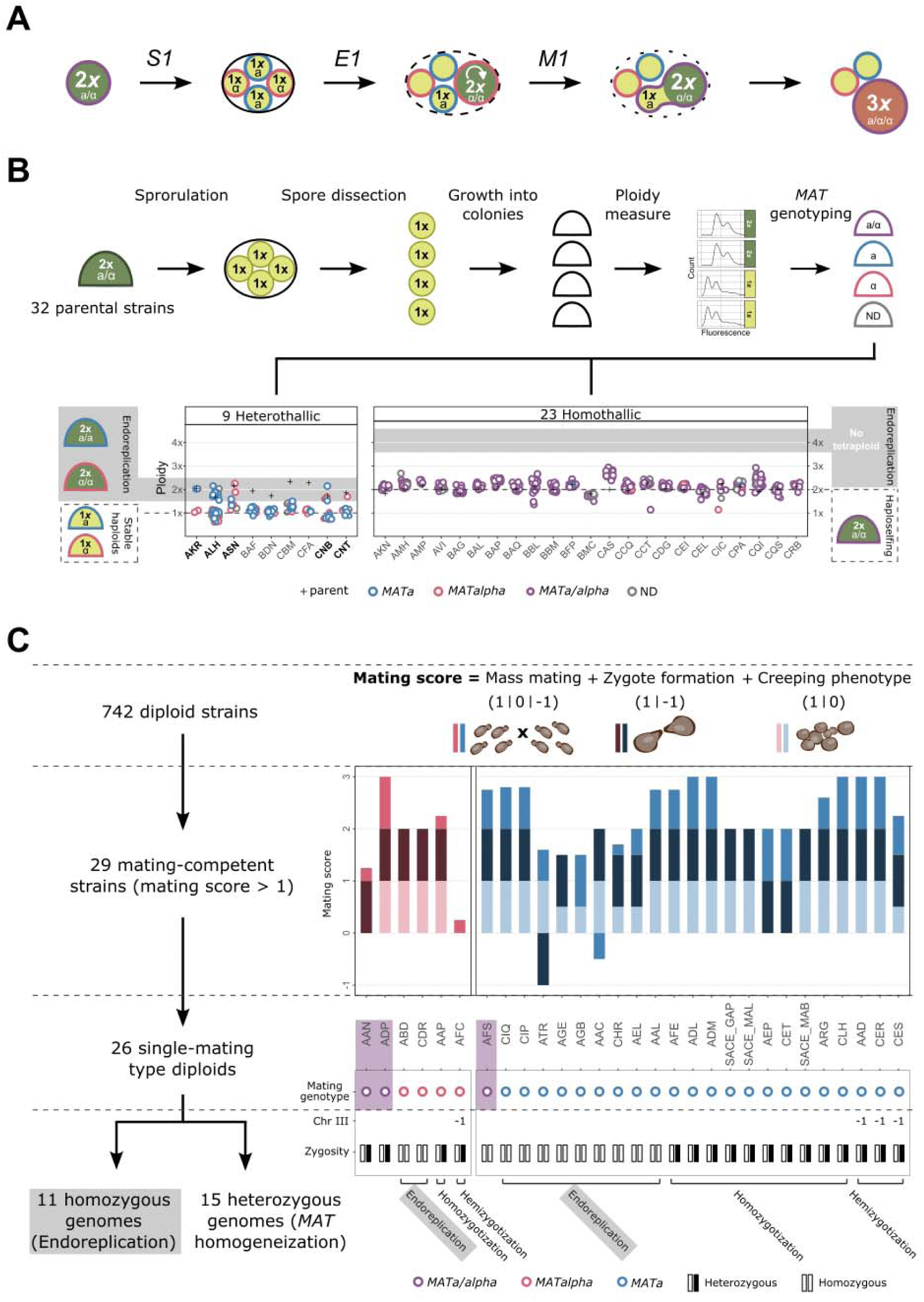
Endoreplication during the first iteration of the *SEM* cycle generates mating- competent diploids. **A**, Schematic of the first iteration of the *SEM* sequence, in which sporulation (*S1*), endoreplication (*E1*), and mating (*M1*) collectively enable the transition from diploid to triploid. Haploid, diploid, and triploid cells or spores are represented by yellow, green, and orange circles, respectively. Circle borders indicate mating type: blue for *MATa*, pink for *MAT*α and purple for *MATa/MAT*α. **B,** Ploidy measurements and mating-type genotyping of colonies derived from individual spores of 23 homothallic (self-mating) and 9 heterothallic (non-self-mating) diploid strains. Individual colonies are represented by semicircles, using the same mating-type color coding as in A. Homothallic strains predominantly produced diploid *MATa/MAT*α colonies (purple) following self-mating, whereas heterothallic strains produced stable haploid *MATa* (blue) or *MAT*α (pink) colonies. Five heterothallic strains (AKR, ALH, ASN, CNB, and CNT) also produced colonies with ploidy values consistent with endoreplication (gray-shaded area). None of the homothallic spore-derived colonies reached the shaded range expected for diploid-to-tetraploid endoreplication. CAS produced colonies with increased ploidy relative to the parental strain (black cross), although below the range expected for endoreplication. **C,** Identification of natural diploid strains capable of mating. Among 742 natural diploid strains screened, 29 showed reproducible mating activity, of which 26 displayed a single mating type and are characterized here. Mating scores combine mass-mating assays (medium shading), direct observation of zygote formation (dark shading), and the creeping phenotype (light shading), a mating-associated aggregation phenotype used as an additional readout of mating competence (Arras *et al*, 2022). Non-integer scores result from averaging replicate measurements across the three assays (see Methods). *MAT* locus homogenization can result either from the loss of one mating-type allele (hemizygotization), resulting from the loss of one copy of chromosome III, or from conversion of the *MAT* locus (homozygotization). Fully homozygous diploids carrying a single mating type are consistent with an origin through haploid endoreplication.

To test this possibility, we screened 32 diploid yeast strains with well-characterized differences in their ability to complete the sexual cycle by self-mating when spores are isolated. We sporulated all strains, isolated 684 spores by microdissection, and determined the ploidy and mating type of the resulting colonies (Table S2, Table S3). Notably, five strains produced 7 to 60% of spore-derived colonies with doubled genome content, showing that endoreplication can occur after spore germination and supporting the existence of the *E1* step of the *SEM* sequence (Fig. 2A).

More specifically, these five strains were heterothallic, meaning that they had lost the ability to switch mating type during the haploid phase and self-mate. Upon sporulation, non-self-mating strains (heterothallic) produce haploid spores that normally develop into single-mating-type haploid colonies. Consistent with this expectation, all nine non-self mating strains tested produced such stable haploid colonies, yet consistent with postmeiotic endoreplication, five also generated apparently diploid colonies, (Fig. 2B). By contrast, homothallic strains produce haploid spores that can switch mating type after germination and self-mate, generating *MATa/MAT*α cells that develop into diploid colonies. Accordingly, spore microdissection showed that all 23 self-mating strains tested produced *MATa/MAT*α diploid colonies, and none showed evidence of endoreplication, which would instead have generated tetraploid colonies (Fig. 2B).

We next tested whether the putatively endoreplicated colonies isolated from the five non-self-mating strains could mate with compatible haploid tester strains and generate triploid progeny, corresponding to the *M1* step of the *SEM* sequence (Fig. 2A). Across the five strains, only the AKR and ALH ∼2*x* spore-derived isolates, both belonging to the Asian fermentation clade, produced triploid progeny upon mating (Supplementary Fig. 1). These results confirm that postmeiotic endoreplication can occur after spore germination and produce mating-competent cells.

The precise timing of endoreplication remains uncertain: it may occur shortly after germination, during the first cell division, or later during colony development. Nevertheless, flow-cytometric profiles of the endoreplicated colonies showed only the characteristic 2× peaks, with no detectable evidence of ploidy heterogeneity. To further assess this homogeneity, we subcloned one endoreplicated colony and measured the ploidy of 10 independent subclones. All subclones displayed a diploid profile (Supplementary Fig. 2), indicating that these colonies were largely homogeneous in ploidy and suggesting that endoreplication occurs early after spore germination, possibly at the first cell division.

We then screened a collection of 742 natural diploid strains from the 1,011 collection using a combination of mating assays (Extended Methods) to identify those that may have arisen through endoreplication of an originally haploid cell without completing the *SEM* sequence. As expected, most strains exhibited no or extremely low mating activity. Nevertheless, a small subset of strains showed clear and reproducible mating signals, including 26 carrying a single mating type and three carrying both mating types (Table S1, Fig. 2C). Of the 26 single-mating-type strains, eleven were fully homozygous across their genomes (Peter *et al*, 2018), indicating that they most likely resulted from endoreplication of an originally haploid cell (Fig. 2C). These strains belonged to eight different clades, suggesting that endoreplication occurs across diverse *S. cerevisiae* populations (Table S1). In addition, we identified 15 single-mating-type diploids capable of mating that were heterozygous at the genome level, and therefore not compatible with an endoreplication origin. Their mating ability likely arose through homogeneization of their *MAT* locus (Fig. 2C). These strains could contribute to polyploidization through outcrossing, independently from the *SEM* sequence. This screen provides clear evidence that endoreplication spontaneously occurs at detectable frequencies and that mating-competent heterozygous and homozygous diploids are present in natural yeast populations.

Together, these results demonstrate that (i) postmeiotic endoreplication can occur upon spore germination, (ii) endoreplicated isolates remain capable of mating and generating triploid progeny, and (iii) a subset of natural diploid isolates likely originated through haploid endoreplication. Additionally, these findings are consistent with the previously reported association between loss of self-mating ability and natural polyploidy (Vittorelli *et al*, 2026), as only heterothallic diploids generated triploid progeny through the *SEM* pathway in our experiments.

### Second *SEM* cycle: stepwise tetraploidization

The second iteration of the *SEM* cycle drives the stepwise diploid-to-triploid transition. It starts with *S2*, sporulation of a triploid parental cell (Fig. 3A). Among 46 triploids from the 1,011 collection, 38 (82.6%) were able to sporulate, with efficiency ranging from 3% to 97% (median 44%) (De Chiara *et al*, 2022). To assess triploid fertility, we microdissected 180 spores from each of three representative euploid triploid strains and determine their viability. Spore viability was extremely low, ranging from 5.0–7.8% (Supplementary Fig. 3, Table S3). Thus, although natural triploids frequently undergo sporulation, the resulting spores have very low viability.

**Figure 3:**
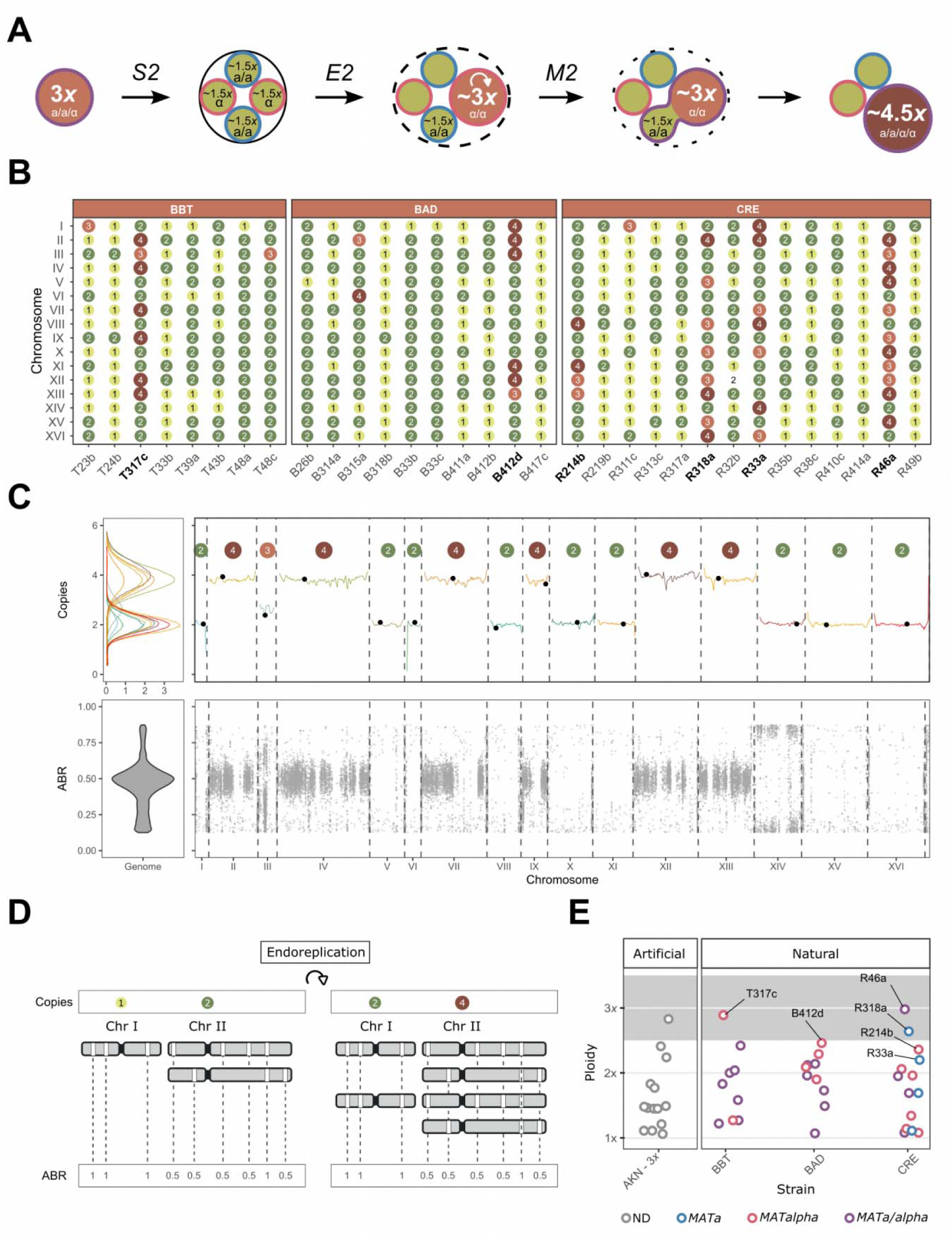
The second iteration of the *SEM* cycle enables stepwise triploid-to-tetraploid transition. **A**, Schematic of the second *SEM* iteration (*S2–E2–M2*) initiated by sporulation of a triploid parental strain. Triploid meiosis produces highly aneuploid spores with variable chromosome copy numbers that average 1.5*x*. The mating genotype of the parental triploid strain exemplified here is *MATa/*α*/*α. **B**, Chromosome copy-number profiles of 32 colonies derived from individual spores following meiosis of three triploid parental strains (BBT, BAD, and CRE), showing extensive aneuploidy. Six isolates (names in bold) contain chromosomes present in four copies and completely lack single- copy chromosomes, two signatures consistent with postmeiotic endoreplication. **C**, Genome-wide (left) and chromosome-level (right) profiles of chromosome copy number (top) and allele balance ratio (*ABR*; bottom) for the T317c spore-derived isolate. Chromosome copy number was estimated from sequencing depth normalized to genome ploidy. Centromeres are marked with black circles, and integer chromosome copy numbers are indicated in colored circles. In the chromosome-level *ABR* profile, each gray point represents a heterozygous variant along the chromosome. *ABR* profiles show the fraction of sequencing reads carrying the alternative allele at heterozygous sites, only variants with *ABR* values between 0.125 and 0.875 are shown (see Methods). **D**, Schematic illustrating the effect of endoreplication on copy number and allele balance ratio (*ABR*), using Chr I and Chr II as examples. Following genome doubling, a chromosome initially present in one copy becomes a homozygous two-copy chromosome, with inherited variants displaying an *ABR* of 1. In contrast, an initially heterozygous two-copy chromosome becomes four-copy, with heterozygous variants retaining an *ABR* of ∼0.5. **E**, Estimated ploidy of the 32 spore-derived isolates. The six putatively endoreplicated isolates identified in panel B are highlighted and show increased ploidy relative to the remaining isolates. The shaded area indicates the approximate ploidy range expected following endoreplication.

To quantify this effect, we constructed a fully homozygous triploid strain (AKN-3*x*) and measured its spore viability. This artificial triploid showed 56.7% spore viability (Supplementary Fig. 3), compared with 96% in the original AKN diploid strain (Table S3), indicating that segregation defects alone can reduce fertility by approximately twofold in triploids, as previously reported (Charles *et al*, 2010). The substantially lower spore viability of natural triploids (5.0–7.8%) suggests that, in addition to triploidy-associated segregation defects, meiotic segregation unmasks a major contribution from recessive deleterious variants in their genomes, including several heterozygous high-impact variants in essential genes (Mozzachiodi *et al*, 2022).

Because triploid cells carry an uneven number of homologouss chromosome, these copies cannot be evenly segregated during meiosis. As a result, the spores produced during the *S2* step inherit variable numbers of chromosomes, ranging from 1*x* to 3*x* with an expected average of 1.5*x* (Fig. 3A). We sequenced 32 colonies derived from individual spores of three natural triploids. All 32 colonies were aneuploid, carrying unequal numbers of different chromosomes rather than complete haploid, diploid or triploid chromosome sets (Fig. 3B; Table S2). Six of the 32 sequenced isolates showed four genomic signatures consistent with *E2* (Fig. 3A; Supplementary Fig. 4). First, these isolates contained several chromosomes present in four copies (Fig. 3B). Because triploid meiosis can generate at most three copies of any chromosome in a spore, four-copy chromosomes are most parsimoniously explained by duplication of chromosomes initially present in two copies. Second, none of these six isolates contained single-copy chromosomes, which were common among the remaining isolates. This absence is also consistent with endoreplication, as chromosomes initially present in a single copy would be duplicated to two copies (Fig. 3B). Third, allele balance ratios (*ABRs*), which measure the relative abundance of alternative alleles at heterozygous sites, showed patterns consistent with genome doubling (Fig. 3C). In four-copy chromosomes, heterozygous variants had an *ABR* of ∼ 0.5, as expected when two-copy chromosomes are duplicated (Fig. 3D). By contrast, two-copy chromosomes lacked heterozygous variants, consistent with duplication of chromosomes that were initially present in a single copy (Fig. 3C and 3D; Supplementary Fig. 4). Finally, consistent with genome doubling, the six putatively endoreplicated isolates also had higher overall ploidy (2.2*x*–3.0*x*; mean 2.6*x*) than the remaining isolates (1.1*x*–2.4*x*; mean 1.7*x*; Fig. 3E). Together, these genomic signatures indicate that 6 of 32 sequenced triploid-derived isolates (19%) underwent postmeiotic endoreplication, supporting the *E2* step of the *SEM* sequence (Fig. 3A).

We further asked whether the six putatively endoreplicated spore-derived isolates retained mating competence, thereby testing the *M2* step of the *SEM* sequence (Fig. 3A). We determined their mating types and assessed their ability to mate in mass mating assays, using 26 non-endoreplicated isolates for comparison. Five of the six putatively endoreplicated isolates were *MAT*_α_ and mating-competent, whereas one was unable to mate (Table S2). We then measured the ploidy of the resulting progeny. Four of the five mating-competent endoreplicated isolates produced progeny with increased ploidy, with gains ranging from +0.7*x* to +2.1*x* relative to the parental isolates (Supplementary Fig. 5). Thus, five out of six endoreplicated triploid-derived spores retained mating competence and could further increase their ploidy through mating, supporting the *M2* step of the *SEM* sequence (Fig. 3A).

Together, these results show that triploids frequently undergo sporulation (*S2*) but produce few viable spores. Among those a significant subset undergoes postmeiotic endoreplication (*E2*), wile retaining the ability to mate (*M2*). Thus, despite their low fertility, triploids can complete a second *SEM* cycle, enabling a stepwise increase in ploidy toward tetraploidy.

### Spontaneous generation of novel polyploids through *SEM*

A key prediction of the *SEM* sequence is that postmeiotic endoreplication and mating occur sequentially after spore germination while the spores remain in close physical proximity following rupture of the ascus. To directly test this prediction, we individually transferred 1,191 intact asci from 12 non-self-mating diploid strains onto rich medium, in contrast to the previous experiments in which individual spores were isolated. These strains represented diverse genetic and ecological backgrounds (Table S2). We focused on heterothallic strains, which cannot switch mating type during their haploid phase, because our previous experiments showed that spore endoreplication occurred only in these backgrounds (Fig. 2C) and because the loss of selfing ability is strongly associated with natural polyploidy (34). We allowed each ascus to develop into a colony, recorded ascus viability, and determined the ploidy of the resulting colony (Fig. 4A, Table S4).

**Figure 4:**
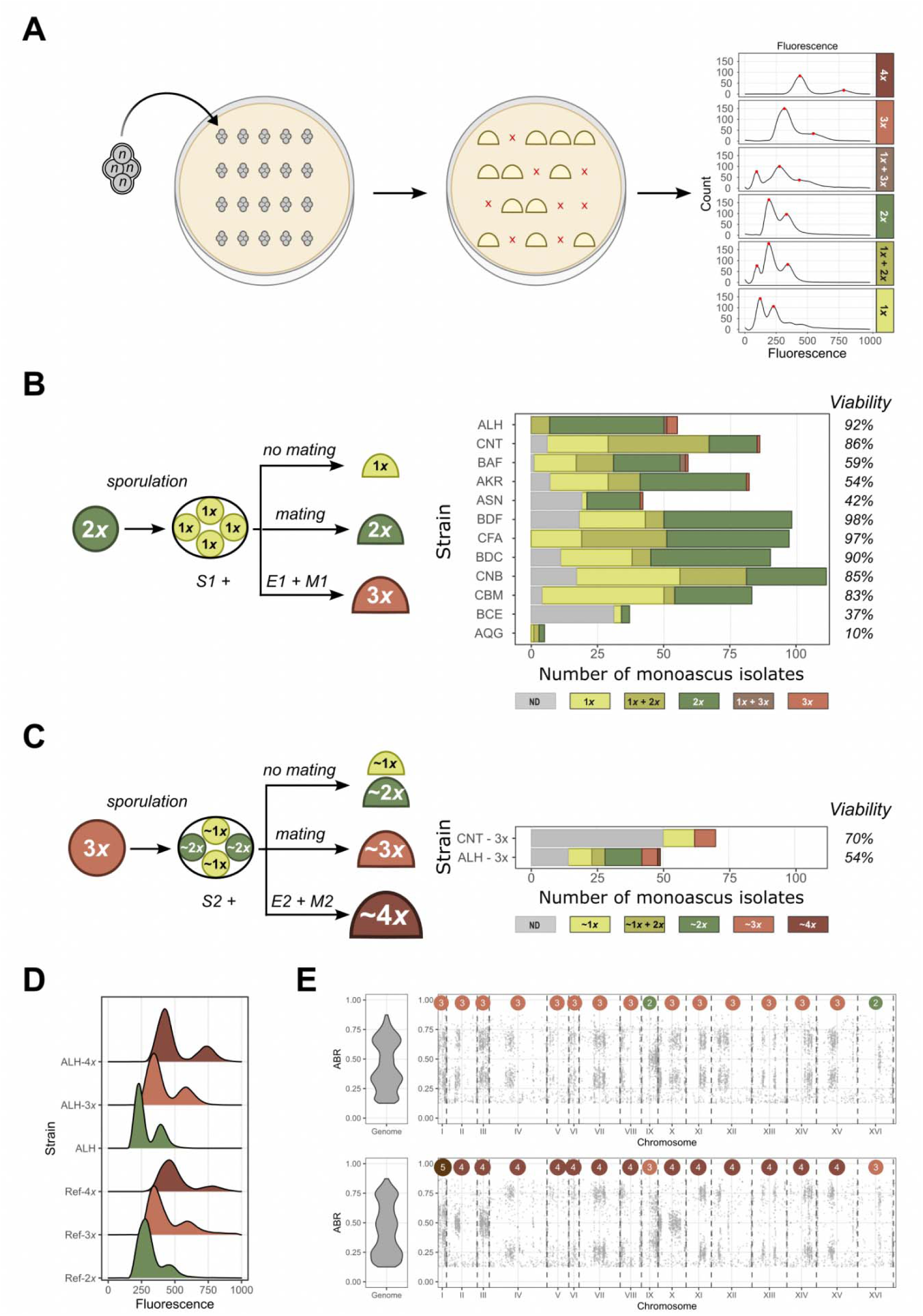
Spontaneous generation of novel polyploids through iterative *SEM* cycles. **A**, Experimental design using intact asci, allowing spores within the same ascus to germinate, undergo potential endoreplication, and mate before developing into colonies. The ploidy of viable colonies was determined by flow cytometry. Histograms on the right show DNA-content measurements from representative colonies, and are not schematic. **B**, Ploidy outcomes of colonies derived from individual asci of heterothallic diploid strains. Triploid isolates (3*x*) arise spontaneously in multiple genetic backgrounds, consistent with completion of the first *SEM* cycle. Some colonies contained cells with mixed ploidies, including haploid and diploid (1*x*+2*x*), or haploid and triploid (1*x*+3*x*) cells. In other cases, ploidy could not be reliably determined (ND) due to slow growth or multicellular clumping, a phenotype frequently observed in wild strains (Barrere *et al*, 2023). **C**, Ploidy outcomes following sporulation of newly formed triploids. The resulting colonies span a broad range of ploidies, including nearly tetraploidy (∼4*x*), consistent with completion of the second *SEM* cycle. **D**, DNA-content profiles by flow cytometry for ALH-derived isolates of different ploidies, including ALH-2*x*, ALH-3*x*, and ALH-4*x*. Reference strains used for ploidy calibration were the laboratory strain BY4743 (2*x*), the natural triploid CRE (3*x*) and the artificial tetraploid AKN-4*x*. **E**, Genome-wide (left) and chromosome-level (right) allele balance ratio (*ABR*) profiles of ALH-3*x* (top) and ALH-4*x* (bottom). Colored circles indicate chromosome copy numbers estimated from sequencing depth normalized to genome ploidy. *ABR* profiles show the fraction of sequencing reads carrying the alternative allele at heterozygous sites (y-axis), with each gray point representing a variant along the chromosome (x-axis). In ALH-3*x*, *ABRs* cluster around ∼0.33 and ∼0.67 on three-copy chromosomes, corresponding to 1:2 and 2:1 allelic ratio, respectively. In ALH-4*x*, *ABRs* cluster around ∼0.25, ∼0.5, and ∼0.75 on four-copy chromosomes, corresponding to 1:3, 2:2, and 3:1 allelic ratio, respectively.

Ascus viability varied widely among strains (Fig. 4B), and the resulting colonies were predominantly haploid or diploid. Colonies remained haploid when no mating occurred after germination, whereas mating between 1*x* compatible spores from the same ascus resulted in diploid colonies (Fig. 4B). Additionally, five of the twelve tested strains also produced triploid colonies, at frequencies ranging from 1.2% to 9.1% (Fig. 4B), consistent with completion of a first *SEM* cycle, in which one 1*x* spore first undergoes endoreplication to become 2*x* and subsequently mates with another 1*x* spore from the same ascus (Fig. 2A).

In total, eight independent triploid isolates were recovered, subcloned, and confirmed as triploid by both flow cytometry and whole-genome sequencing (Tables S4 and S5). Their chromosome copy-number and *ABRs* showed the expected signatures of triploidy (Supplementary Fig. 6). Together, these results demonstrate that sporulation of non-self-mating diploids can spontaneously generate triploid progeny through completion of a *SEM* cycle (Fig. 2A).

We further tested whether the newly formed triploids could undergo a second *SEM* cycle and generate tetraploid progeny. We selected one triploid isolate from each of two genetic backgrounds (ALH and CNT) and induced sporulation again. In both cases, ascus viability was lower than in the corresponding diploid parent, consistent with the reduced fertility of triploids (Fig. 4C). The resulting colonies displayed a broad range of ploidies, from near-haploid to near-tetraploid. Near-haploid and diploid colonies likely resulted from uneven chromosome segregation, a characteristic of triploid meiosis in the absence of mating, whereas near-triploid colonies are consistent with ploidy restoration through mating between spores within the same ascus (Fig. 4C). Notably, we identified a tetraploid isolate, ALH-4*x,* among the progeny of ALH-3*x* triploid isolate (Fig. 4C), consistent with completion of a second *SEM* cycle, in which an approximately 1.5*x* spore undergoes endoreplication to reach ∼3*x* and subsequently mates with a near-haploid spore from the same ascus (Fig. 3A).

To validate this tetraploid transition, ALH-4*x* was subcloned and its ploidy was confirmed by both flow cytometry and whole-genome sequencing (Fig. 4D and E). Comparison of chromosome copy-number and *ABRs* between ALH-3*x* and ALH-4*x* revealed the expected +1*x* increase in genome content, providing direct evidence that a second *SEM* cycle can increase ploidy from triploidy to tetraploidy (Fig. 3A).

Together, these results demonstrate that the stepwise diploid-to-tetraploid transition can proceed through a triploid intermediate via two successive *SEM* cycles under laboratory conditions.

### A barrier to increasing ploidy beyond tetraploidy

We next asked whether additional *SEM* cycles could drive further increases in ploidy beyond tetraploidy. Among 38 natural tetraploids, 25 (65.8%) were able to sporulate, with a median sporulation efficiency of 57% (De Chiara *et al*, 2022). However, analysis of the meiotic progeny of two representative tetraploid strains revealed low spore viability (6.7%–12.2%, Supplementary Fig. 3). In addition, we detected no evidence of spore endoreplication in tetraploid-derived progeny, suggesting that further ploidy increases through the *SEM* sequence are limited, or at least less frequent than in triploid progeny (Supplementary Fig. 7A).

More specifically, genome sequencing of 28 colonies derived from individual spores of two natural complete tetraploids revealed relatively few chromosome-number abnormalities compared with triploid progeny (Supplementary Fig. 7B). Eleven isolates were complete diploids, whereas the remaining isolates contained single-copy chromosomes (seven cases), three-copy chromosomes (14 cases), or both. Thus, the low spore viability of natural tetraploids likely reflects a substantial contribution of recessive deleterious variants that become exposed following meiotic segregation. To test this interpretation, we constructed an artificial homozygous tetraploid (AKN-4*x*) to estimate spore viability in the absence of recessive deleterious variants. This strain showed 77.2% spore viability, substantially higher than the 56.7% in the homozygous triploid AKN-3*x* (Supplementary Fig. 3), consistent with chromosome segregation being less affected in tetraploids than in triploids, as previously reported (Loidl, 1995).

We extended this analysis to a natural pentaploid. Genome sequencing of 16 colonies derived from individual spores revealed predominantly chromosomes present in two or three copies, with only a single chromosome present in one copy across all isolates (Supplementary Fig. 7B). As in tetraploids, we detected no evidence of spore endoreplication.

In conclusion, we found no evidence for stepwise ploidy increases beyond tetraploidy, suggesting that 4*x* might represent a natural barrier for *SEM*-mediated polyploidization. This is consistent with the rarity of natural pentaploids, which account for only about 0.2% of strains in the largest available strain collection (Loegler *et al*, 2024), and with the absence of hexaploid or higher-ploidy isolates.

### Stepwise polyploidization as a major route to natural polyploidy

Because saltational and stepwise polyploidization arise through different mechanisms (Fig. 1), they are expected to leave distinct genomic signatures. To distinguish between these two routes, we analyzed the genomes of 91 natural polyploids from the 1,011 collection (Peter *et al*, 2018) and asked whether their genomic features were more consistent with a stepwise or a saltational origin.

We first reassessed the DNA content of the 91 natural polyploids by flow cytometry to infer ploidy and analyzed their genome-wide allele balance (Fig. 5A). The data revealed an almost continuous ploidy range from 2.5x to 5.5x (Fig. 5A, Table S6), with clear discontinuities near 3.5x and 4.5x separating triploid, tetraploid, and pentaploid classes. After reassignment of a few strains, the final dataset comprised 48 triploids, 38 tetraploids, and 5 pentaploids (Table S6). The prevalence triploids is difficult to reconcile with a saltational polyploidization model (Fig. 1A). In plants, triploids can arise through fusion of an unreduced gamete generated by meiotic restitution with a fully reduced gamete from a separate meiosis. This route to triploidy may be more constrained in yeast because (i) skipping the first meiotic division would produce two *MATa/MAT*_α_ diploid cells unable to mate and (ii) second-division restitution would produce two unreduced spores within the same ascus that can mate to generate tetraploids. Thus, under the saltational model, triploid yeast cells would instead arise through chromosome loss from tetraploids. Yet, although rapid chromosome loss has been documented in laboratory tetraploids (Mayer & Aguilera, 1990; Gerstein *et al*, 2006, 2008), it typically produces diploid rather than triploid genomes (Selmecki *et al*, 2015). Moreover, long-term vegetative propagation of natural tetraploids revealed that ploidy levels remain stable (Dutta *et al*, 2022). Together these observations argue against chromosome loss from tetraploids as a major route to triploid formation.

**Figure 5:**
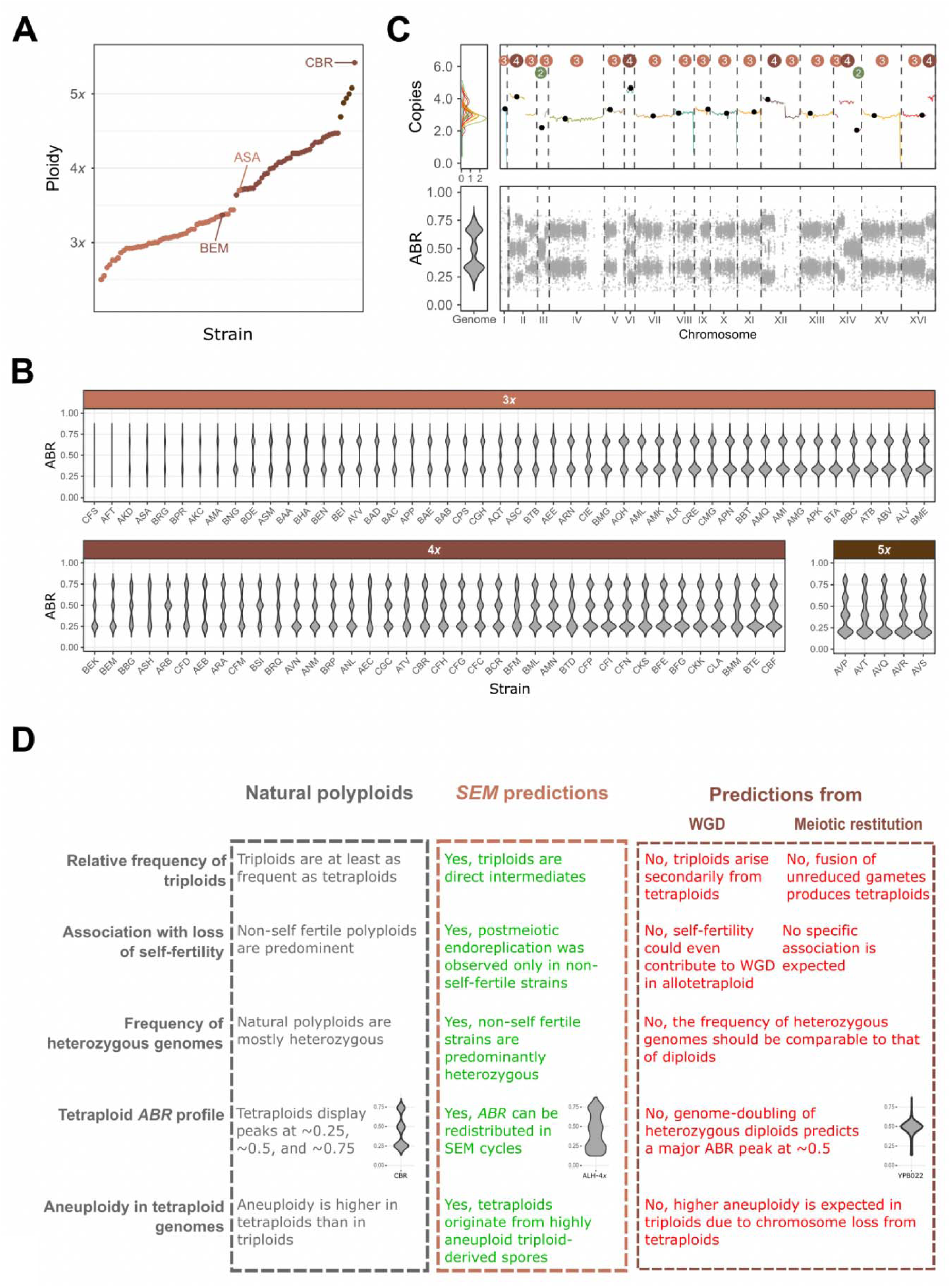
Genomic signatures of natural polyploids support a stepwise origin. **A**, Ploidy estimates of 91 natural polyploid isolates inferred from flow cytometry, showing a near-continuous distribution from ∼2.5*x* to ∼5.5*x* with discontinuities delineating triploid, tetraploid, and pentaploid classes. Genome-wide allele-balance estimates were largely concordant with these measurements; BEM, ASA, and CBR are highlighted as the three exceptions. **B**, Genome-wide allele balance ratio (*ABR*) distributions for the 91 natural polyploids. The observed *ABR* patterns are inconsistent with whole-genome duplication of heterozygous diploids and instead support stepwise increase in ploidy. **C**, Chromosome copy-number and *ABR* profiles of a representative natural polyploid strain (AQT) presenting several whole-chromosome and segmental aneuploidies. **D**, Summary of the genomic signatures observed in natural polyploids and their agreement or disagreement with predictions of the stepwise and saltational routes to polyploidization. Examples of *ABR* profiles illustrate different peak distributions observed in tetraploids. CBR represents a natural tetraploid, ALH-4*x* a tetraploid generated through two successive *SEM* cycles, and YPB022 a laboratory-generated tetraploid produced by combining the YPS128 and YJM1248 strains from the 1,011 collection.

The prevalence of strains unable to switch mating type, and therefore unable to self-mate, among polyploids is evident in the 1,011 collection: among 91 polyploids, 73 were unable to self-mate, whereas only 7 were self-mating and 11 were of unknown status (Table S6) (Vittorelli *et al*, 2026). Consistent with this pattern, postmeiotic endoreplication during the *SEM* cycle was observed only in non-self-mating isolates in our experiments. By contrast, the saltational model does not predict a specific association between loss of self-fertility and polyploidization. Indeed, mating-type switching may even facilitate diploid-to-tetraploid transitions involving whole-genome duplication, as suggested for the formation of allotetraploids (Ortiz-Merino *et al*, 2017). The strong enrichment of strains unable to self-mate among natural polyploids is therefore consistent with, and provides additional support for, a stepwise route to polyploidization.

Overall patterns of genome homozygosity and heterozygosity were also inconsistent with a predominantly saltational origin. If WGD occurred independently of whether the parental genome was homozygous or heterozygous, the fraction of homozygous polyploids should be comparable to that observed among diploids (47% in the 1,011 collection; (Peter *et al*, 2018)). Instead, we detected only two homozygous triploids and no homozygous tetraploids among the 91 natural polyploids. Together, the association of polyploidy with loss of self-fertility and elevated genome-wide heterozygosity further supports a stepwise origin of natural polyploids.

Furthermore, WGD of a heterozygous diploid genome would produce tetraploids in which heterozygous variants are present in approximately 50% of sequencing reads, resulting in a major peak at an *ABR* of ∼0.5 (Supplementary Fig. 8). By contrast, all 38 natural tetraploids displayed three distinct peaks at *ABR* ∼0.25, ∼0.5, and ∼0.75 (Fig. 5B), a pattern also observed in the ALH-4*x* isolate generated through two successive *SEM* cycles (Fig. 4E). The additional peaks at ∼0.25 and ∼0.75 are inconsistent with simple WGD of a heterozygous diploid and instead supports a stepwise origin through the *SEM* sequence.

Finally, the saltational route predicts higher aneuploidy in triploids arising through chromosome loss from tetraploids (Gerstein *et al*, 2008), whereas the *SEM* model predicts higher aneuploidy in tetraploids, which originate from highly aneuploid triploid-derived spores. Natural polyploid genomes show extensive aneuploidy in both triploids and tetraploids, including 122 whole-chromosome and 63 segmental aneuploidies (Table S6, Fig. 5C). Tetraploids contain a higher proportion of aneuploid strains than triploids (78% versus 52%), a pattern consistent with the *SEM* model and with the higher chromosome-loss rate reported for tetraploids than for triploids (Mayer & Aguilera, 1990).

Together, these genomic signatures argue against a dominant saltational origin and support a stepwise polyploidization through iterative *SEM* cycles as a major route to natural polyploidy in *S. cerevisiae* (Fig. 5D).

## Discussion

Here, we identify a novel route to polyploidization in yeast, which we term stepwise polyploidization and propose to be mediated by iterative cycles of the *SEM* sequence. This route begins with sporulation, followed by postmeiotic endoreplication during spore germination, and ends with mating between the endoreplicated spore and a compatible meiotic product that originated from the same ascus and remains nearby. This process differs from the somatic WGD in diploids, which is often invoked to explain polyploid formation in yeast, as well as from the major mechanism described in plants, which relies on the fusion of unreduced gametes produced by meiotic restitution (Ramsey & Schemske, 1998). Notably, a process conceptually related to the *SEM* sequence has been proposed in plants, where a subset of 2*x* nuclei in mature pollen may arise through mitotic endoreplication after completion of meiosis rather than through meiotic restitution (Pan *et al*, 2004). This conceptual parallel raises the possibility that postmeiotic endoreplication may contribute to polyploid formation beyond fungi.

Other mechanisms can also generate mating-competent diploids independently of *SEM*. These include *MAT* locus homogenization through unprogrammed mating-type switching or return to growth following aborted meiosis, as well as endoreplication during prolonged vegetative growth of haploid cells (Harari *et al*, 2018; Mozzachiodi *et al*, 2021). Such mating-competent diploids could, in principle, generate triploids by mating with compatible haploid cells. A key distinction, however, is that *SEM* couples the formation of a mating-competent endoreplicated spore with the simultaneous availability of a compatible sibling spore from the same ascus, which remains in close physical proximity after ascus breakdown.

The *SEM* model provides a framework for several genomic characteristics of natural yeast polyploids, including their strong association with loss of self-fertility (Vittorelli *et al*, 2026), and suggests two possible mechanisms that could underlie this relationship. First, spores unable to self-mate may remain in a single–mating-type state for longer after germination, thereby providing a window during which endoreplication can occur. By contrast, spores capable of self-mating may rapidly restore diploidy through mating and therefore be less likely to enter the *SEM* pathway. Alternatively, loss of mating-type switching itself could influence endoreplication: if cells attempt to initiate mating-type switching but the process is impaired, this failure could trigger a cellular response that favors genome duplication. Whether either of these mechanisms, or some other mechanism, promotes endoreplication during or shortly after germination remains to be determined.

The postmeiotic endoreplication step may be key for stepwise polyploidization. Although the precise molecular mechanisms underlying spontaneous spore endoreplication remain to be elucidated, an early study identified a mutation in *CDC31*, a gene required for spindle pole body duplication, as a driver of endoreduplication (Schild *et al*, 1981). To date, at least 16 genes have been implicated in ploidy increase in *S. cerevisiae* (Andreychuk *et al*, 2022). These genes encode proteins that primarily regulate spindle pole body duplication, chromosome segregation, cytokinesis and cell-cycle control, thereby highlighting the central role of cell-division and genome-partitioning pathways in the control of ploidy. Moreover, high rate of endoreplication depends on the presence of a functional copy of *SSD1*, a gene that also regulates aneuploidy tolerance in wild yeast (Tung *et al*, 2021; Hose *et al*, 2020). Variation in these and other genes may therefore contribute to differences among strain backgrounds in the frequency and efficiency with which the complete *SEM* sequence occurs. Future work will be required to assess the extent of natural variation in these genes across populations and to dissect the molecular events governing genome doubling associated with spore germination.

Stepwise polyploidization can explain several characteristic features of natural yeast polyploids. First, triploids are not rare anomalies but expected intermediates produced by a single *SEM* cycle. Because meiosis in triploids is intrinsically unbalanced, viable progeny are highly aneuploid, providing an explanation for the extensive aneuploidy observed in tetraploids. The same stepwise process can also generate and maintain heterozygosity, because increases in ploidy occur through mating between genetically distinct meiotic products, producing allele-balance distributions that differ from those expected after duplication of a single diploid genome.

Beyond its evolutionary significance, the *SEM* mechanism has practical implications. Polyploid yeast strains are widely used in fermentation industries, including brewing and baking, and are also frequently isolated from clinical settings (Peter *et al*, 2018; Duan *et al*, 2018; Loegler *et al*, 2024; Ezov *et al*, 2006; Muller & McCusker, 2009; Albertin & Marullo, 2012). The identification of a natural mechanism underlying their formation, together with our ability to experimentally recapitulate *SEM*-mediated polyploidization under laboratory conditions without genetic modification, opens new avenues for the rational generation of polyploid strains.

In conclusion, our findings support stepwise polyploidization through iterative cycles of the *SEM* sequence as a major route to natural polyploidy in *S. cerevisiae*, while alternative routes, including those involving mating-competent diploids arising from *MAT* hemi- or homozygotization, may also contribute.

## Materials and Methods

### Strains

Natural parental diploid (Table S1) and polyploid (Table S5) strains analyzed in this study were obtained from the 1,011 *S. cerevisiae* collection (Peter *et al*, 2018) and are referred to using the standardized three-letter strain code. Artificial homozygous polyploids derived from the AKN (YPS128) background, as well as heterozygous polyploids derived from AKN and SACE_YBP (YJM1248) backgrounds (YPB012, YPB022, YPB021), were constructed as described previously (Garrido *et al*, 2026). Standard reference strains used for ploidy calibration were as previously described (Gómez-Muñoz & Fischer, 2025). Tester strains used for mass mating assays were OS104a and OS104_α_ (YPS128 background)(Cubillos *et al*, 2009).

### Media composition and growth conditions

Standard growth medium was YPD, consisting of yeast extract (10 g L^-1^), peptone (20 g L^-1^), and dextrose (20 g L^-1^), prepared with or without agar (20 g L^-1^). Sporulation medium was potassium acetate (KAc) agar, containing potassium acetate (20 g L^-1^) and agar (25 g L^-1^). Selective synthetic complete media were prepared using yeast nitrogen base without ammonium sulfate (YNB; 1.7 g L^-1^), arginine (2 g L^-1^), glucose (20 g L^-1^), and agar (20 g L^-1^), supplemented with appropriate complete supplement mixtures (CSM). CSM + G418 medium contained standard CSM and was supplemented with Geneticin (G418; 200 µg mL^-1^). CSM - Ura medium contained CSM lacking uracil (CSM - Ura). CSM - Ura + G418 medium was identical to CSM - Ura and additionally supplemented with G418 (200 µg mL^-1^).

Unless otherwise stated, yeast cells were revived from glycerol stocks on YPD agar and grown at 30 °C until the formation of a visible patch prior to experimental assays. Qualitative sporulation and resporulation capacity were assessed by transferring cells to KAc agar and incubating them at 30 °C. After 7 days, cultures were examined by light microscopy for the presence of asci.

### Chromosome number and ploidy nomenclature

We distinguish chromosome number from ploidy following Ramsey & Schemske (Ramsey & Schemske, 1998). Here, *n* refers to the chromosome number of a gamete and 2*n* to that of a vegetative cell; these terms do not indicate ploidy. Instead, we use *x* to denote ploidy, such that diploid, triploid, and tetraploid cells are 2*x*, 3*x*, and 4*x*, respectively.

### Flow cytometry

Cellular ploidy was estimated by propidium iodide staining followed by flow cytometry. Two sample-preparation procedures were used depending on the purpose of the analysis. For ploidy validations, cells were prepared as previously described (Gómez-Muñoz & Fischer, 2025): cells were grown in YPD, harvested by centrifugation, thoroughly washed with PBS, and stained with propidium iodide (PI; 50 μg/mL). For rapid ploidy screening, samples were prepared either from overnight 1 mL YPD microcultures or by direct fixation of colonies in 70% ethanol. Fluorescence was acquired on a MACSQuant VYB flow cytometer and detected at 605–625 nm, with approximately 10,000 events recorded per sample. Flow cytometry data were gated using FSC-A versus SSC-A and FSC-A versus FSC-H to exclude debris and doublets, followed by FSC-A versus PI fluorescence intensity. When required, cell aggregation was reduced by proteinase K treatment. Ploidy values were inferred using MuPETFlow (Gómez-Muñoz & Fischer, 2025) (see Extended Methods).

### Genotyping mating type loci

Mating genotype was determined by colony PCR as previously described (Vittorelli *et al*, 2026). In a limited number of cases, PCR reactions were performed or repeated using *MAT* locus primers described in (Huxley *et al*, 1990) to confirm mating genotype assignments.

### Mass mating assay

Query and tester cells were mixed on YPD agar and incubated at 30 °C for 24 h to allow mating. Parental query cells were then streaked onto single-selection medium to confirm marker compatibility, while cells from the mating patch were plated onto double-selection medium to assess mating ability (see Media composition and growth conditions).

### Multi-test screening to detect diploid maters

All 742 query strains were initially tested using the mass mating assay with minor modifications, and mating outcomes were scored semi-quantitatively, considering as positive only the crosses that formed a confluent cellular layer (see Extended Methods). A subset of 102 strains was further characterized by microscopic examination of zygote formation after 4 h of co-incubation with tester strains, and by assessment of creeping behavior as previously described (Arras *et al*, 2022) (see Extended Methods). Results from the three assays were integrated into a single composite mating score, and strains with a score greater than 0 were classified as potential maters. Mating efficiency of candidate strains was then evaluated using a quantitative mating assay adapted from (de Barros Lopes *et al*, 2002). Briefly, equal numbers of query and tester cells (2.5 × 10^8^ cells of each) were mixed and spotted onto a nitrocellulose filter placed on YPD agar. After 4 h of co-incubation at 30 °C, cells were recovered from the filter, serially diluted, and plated onto selective media to quantify colony-forming units (CFUs).

Candidate diploid mater strains were subsequently subjected to supplementary analyses to further characterize their mating behavior and genetic features. Mating genotype was determined as described in the section Genotyping mating type loci, and chromosome III aneuploidies and zygosity were retrieved from the original sequencing data (Peter *et al*, 2018).

### Asci dissection and spore isolation

Sporulated asci were treated with lyticase and dissected using a micromanipulator (Singer Instruments MSM 400) as previously described (Vittorelli *et al*, 2026). Spores were plated on YPD agar and cultured at 30 °C, and germination was monitored for up to 7 days. Resulting colonies were picked and stored in glycerol stocks until further analyses.

### Ascus micromanipulation and polyploid detection

Complete asci, irrespective of spore number, were randomly isolated and plated onto YPD agar using a micromanipulator (Singer Instruments MSM 400) and incubated at 30 °C. After colony formation (approximately 2 days), colonies were subjected to rapid ploidy screening by flow cytometry using the procedure described above. Colonies suspected to be polyploid were subsequently single-colony streaked, and their ploidy was confirmed using the validation procedure described above.

### Genomic DNA isolation

Genomic DNA was extracted either using the E-Z 96 Tissue DNA Kit (Omega Bio-Tek), following the manufacturer’s instructions, or using an SDS-potassium acetate (SDS-Kac) protocol, in which cells were lysed with SDS and proteins and cellular debris were precipitated with potassium acetate before DNA recovery, as previously described (Agier *et al*, 2025).

### Genome sequencing and bioinformatic analysis

Genomic DNA was sequenced using Illumina paired-end sequencing (2 × 150 bp) with an average insert size of approximately 300 bp. Read quality was assessed using FastQC (v0.12.1)(Andrews, S. (2010) FastQC A Quality Control Tool for High Throughput Sequence Data. - References - Scientific Research Publishing) and summarized with MultiQC (v1.13)(Ewels *et al*, 2016). Read trimming and filtering were performed using Trimmomatic (v0.39)(Bolger *et al*, 2014), with parallel execution managed using GNU Parallel (v20190322)(GNU Parallel 2018).

Read alignment and variant calling were performed using a custom Snakemake-based pipeline (https://github.com/CintiaG/poly_CCN_and_VC.git). Sequencing reads were aligned to the reference genome using BWA (v0.7.18)(Li & Durbin, 2009). SAM files were converted to BAM format and sorted using SAMtools (v1.21)(Li *et al*, 2009). Duplicate reads were identified and marked using GATK MarkDuplicatesSpark (v4.5.0.0)(Van der Auwera *et al*, 2013). Variants were called per strain using GATK HaplotypeCaller in gVCF mode. Individual gVCF files were merged using GATK CombineGVCFs, and joint genotyping was performed with GATK GenotypeGVCFs to generate final VCF files.

Chromosome copy number (CCN) was estimated from sequencing depth using BEDTools (v2.29.2)(Quinlan & Hall, 2010). CCN and segmental aneuploidies were inferred using in-house scripts (plot_depth.R and plot_copies.R; https://github.com/CintiaG/wgs-aneuploidy-abr.git).

Allele balance ratio (*ABR*) was calculated by extracting read depth (DP) and allele depth (AD) information from VCF files using BCFtools (v1.13)(Danecek & McCarthy, 2017). Text outputs were converted to fst format using the fst package (v0.9.8, http://www.fstpackage.org). *ABR* values were computed using the allele_balance_ratio.R script (https://github.com/CintiaG/wgs-aneuploidy-abr.git), retaining variants with *ABR* between 0.125 and 0.875. The number of heterozygous sites was estimated considering only variants with up to three alleles. In selected cases, CCN estimates were manually refined using *ABR* information, and genome-wide visualizations were generated using the plot_allele_balance_ratio.R script (https://github.com/CintiaG/wgs-aneuploidy-abr.git).

### Data, Materials, and Software Availability

The data for this study have been deposited in the European Nucleotide Archive (ENA) at EMBL-EBI under accession number PRJEB108394. The genomic data analysis pipeline is deposited at the following repository: https://github.com/CintiaG/wgs-aneuploidy-abr.git

## Supporting information

Extented methods

## ACKNOWLEDGMENTS

We are grateful to our colleagues in the *Biology of Genomes* team for their valuable discussions and insights. We thank Nikolas Vakirlis and Fredy Barneche for their critical reading of the manuscript. This work was supported by the Agence Nationale de la Recherche ANR-20-CE12-0020, ANR-24-CE12-0998 and ANR-26-CE12-0292.

