## Supplementary figures for "A postmeiotic route to polyploidy"

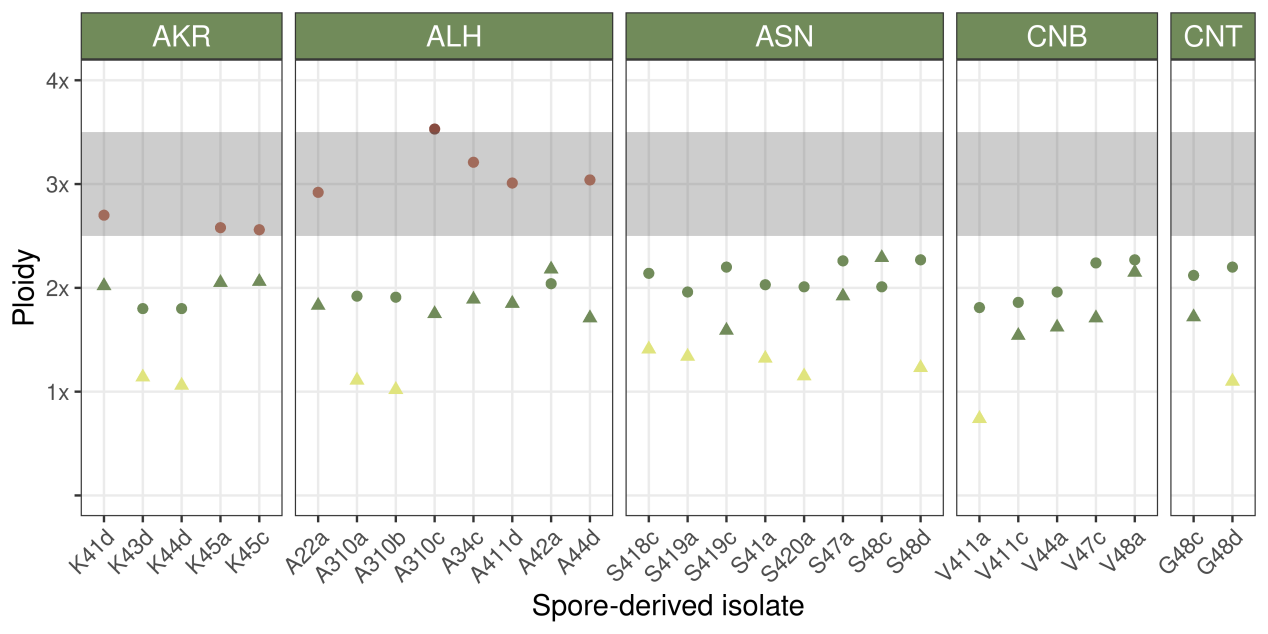
**Figure S1.** Ploidy of spore-derived isolates derived from diploid strains (triangles) and of the progeny generated after crossing these isolates with compatible haploid testers (circles). Each symbol represents a distinct isolate or mating-derived progeny and not a replicate measurement. Ploidy values were estimated from DNA content and therefore may deviate from integer values. Measurements are colored according to the rounded ploidy value: yellow, haploid; green, diploid; orange, triploid; and red, tetraploid. The shaded gray area indicates the expected ploidy range of triploids originating from endoreplicated spores. Spore-derived isolates from strains AKR and ALH produced triploid progeny.

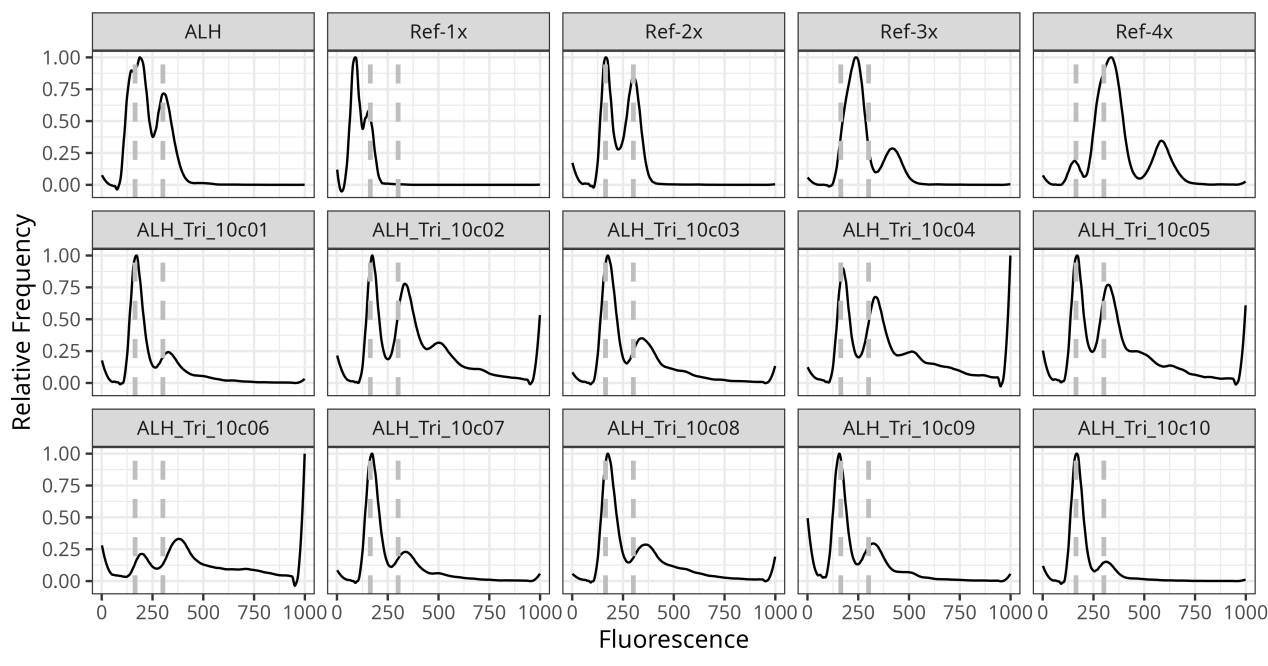

**Figure S2.** Ploidy analysis of subclones derived from an endoreplicated spore-derived colony (ALH_Tri_10c). Flow-cytometric DNA-content profiles are shown for the parental ALH spore-derived colony, haploid (Ref-1x), diploid (Ref-2x), triploid (Ref-3x), and tetraploid (Ref-4x) reference strains, and 10 independent subclones derived from the ALH spore-derived colony. Vertical dashed lines indicate reference fluorescence peak positions for comparison. All 10 subclones displayed DNA-content profiles consistent with diploidy, supporting ploidy homogeneity within the original colony and being consistent with endoreplication occurring early during colony establishment.

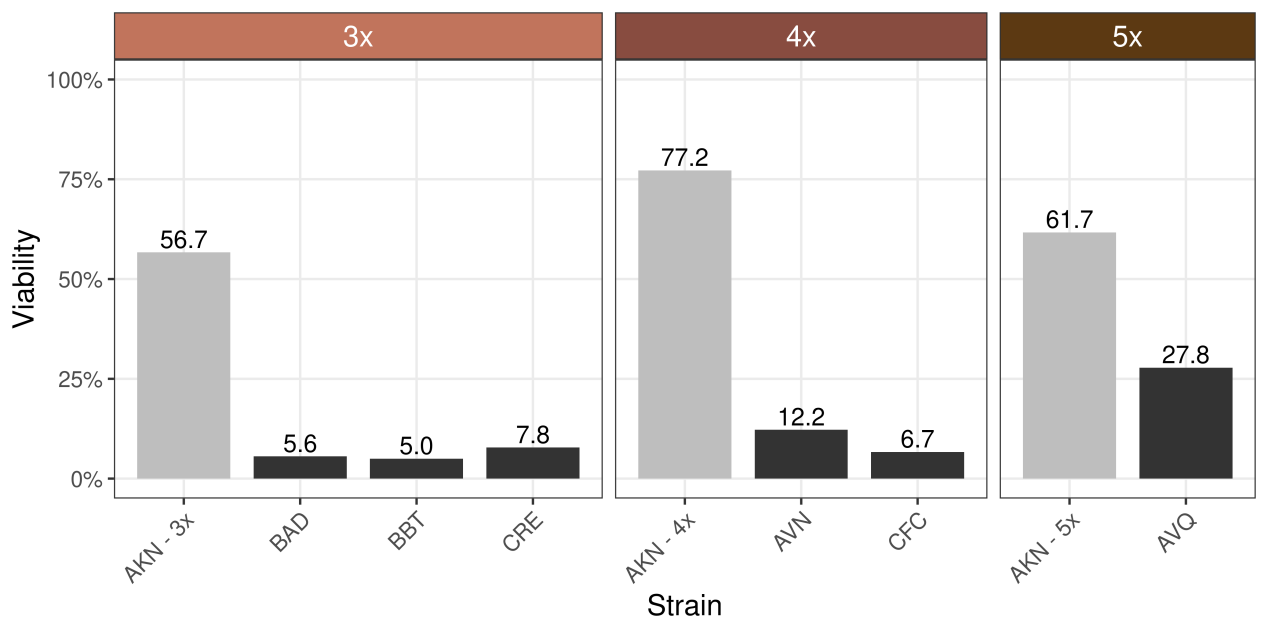

**Figure S3.** Spore viability of artificial homozygous (light gray) and natural heterozygous polyploid strains (dark gray) of varying ploidy. Viability values were inferred from 180 dissected spores per strain. The reduced viability observed in natural strains likely reflects the combined effects of chromosome missegregation and recessive deleterious variants unmasked following meiotic segregation.

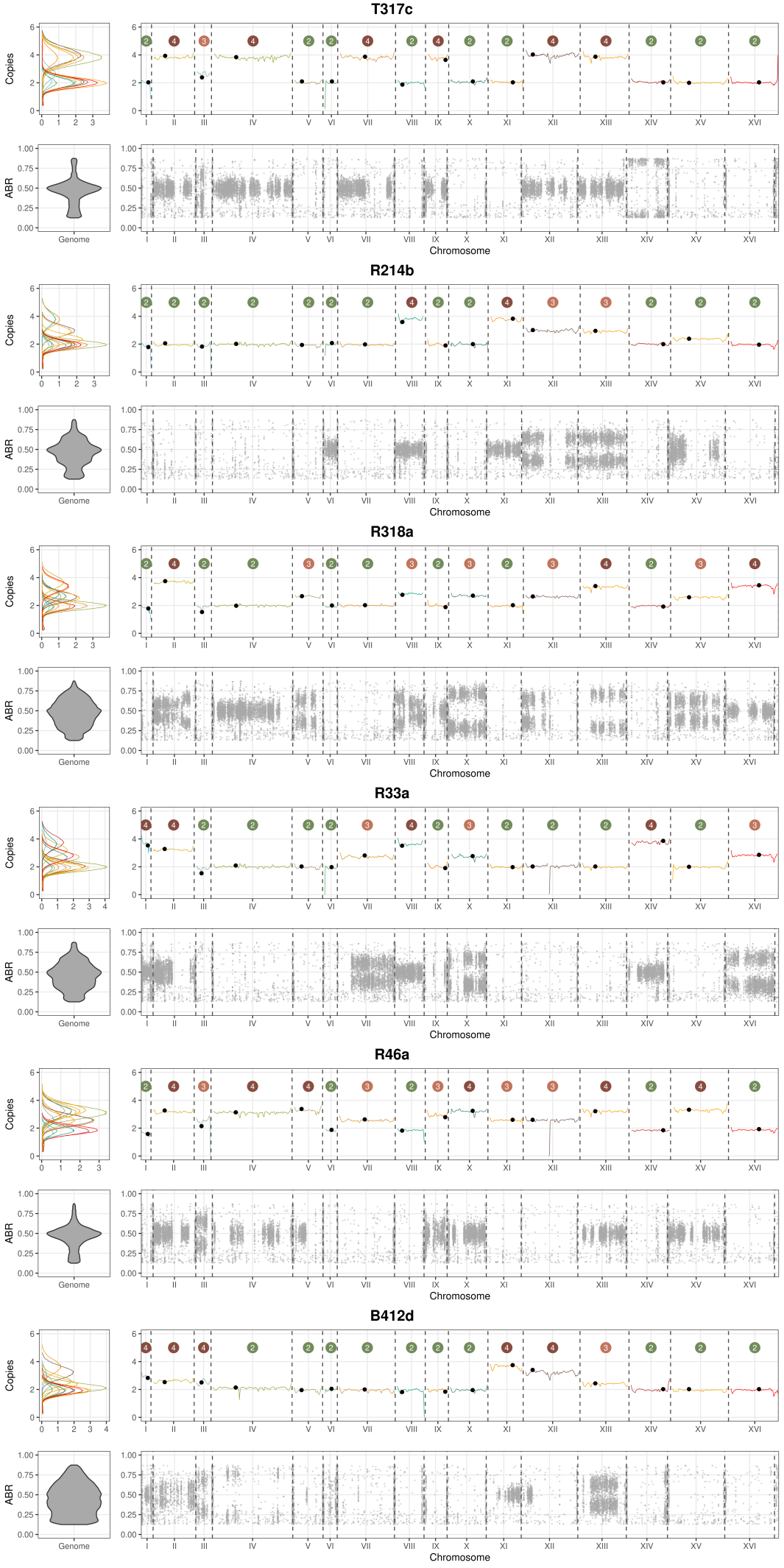

**Figure S4.** Whole-genome profiles of spore-derived isolates showing signatures of postmeiotic endoreplication in triploids strains. Sequencing depth histograms (top left panel) were used to estimate chromosome copy number (top right panel; values indicated above each chromosome). Centromeres are marked with black circles. Allele balance ratio (*ABR*) profiles are shown at the genome-wide level (bottom left) and for individual chromosomes (bottom right), with each point representing an alternative allele at a heterozygous site. Endoreplicated isolates contained homozygous two-copy chromosomes and heterozygous four-copy chromosomes. On four-copy chromosomes, *ABRs* cluster around ~0.5, corresponding to a 2:2 allelic ratio and consistent with duplication of heterozygous two-copy chromosomes during endoreplication.

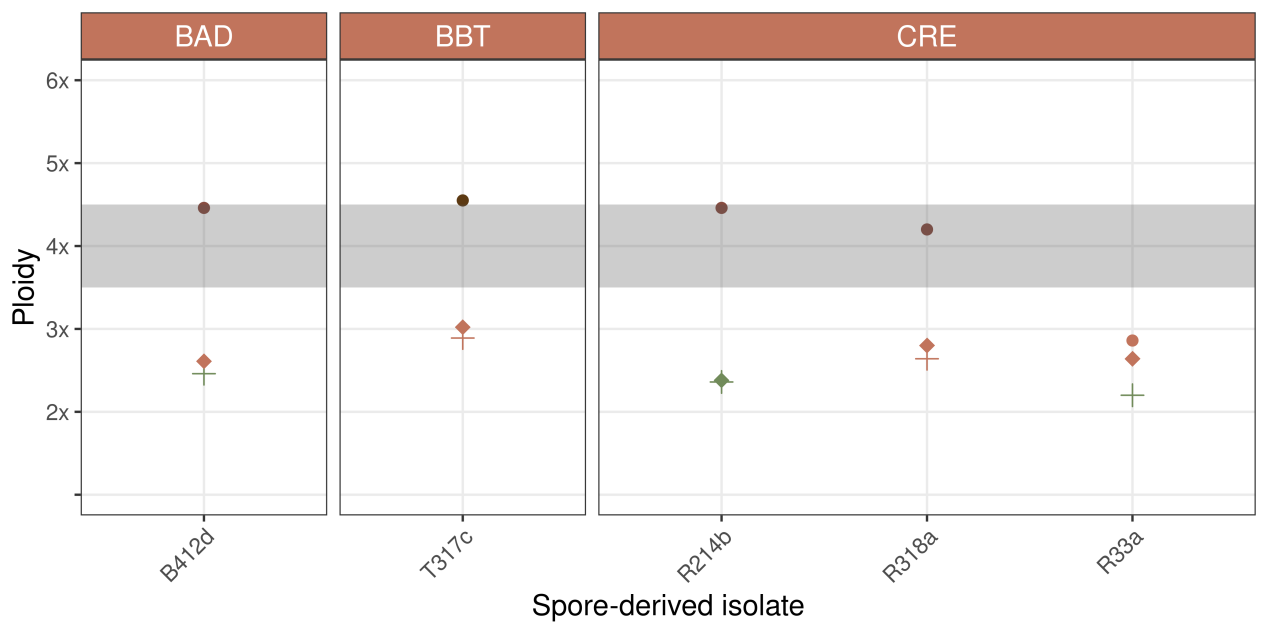

**Figure S5.** Ploidy of spore-derived isolates from triploid strains (crosses), bioinformatically estimated ploidy (diamonds), and their progeny following crosses with compatible haploid testers (circles). Measurements are colored according to the rounded ploidy value: green, diploid; orange, triploid; red, tetraploid; and dark red, pentaploid. The shaded gray area highlights the approximate ploidy threshold observed for the crosses.

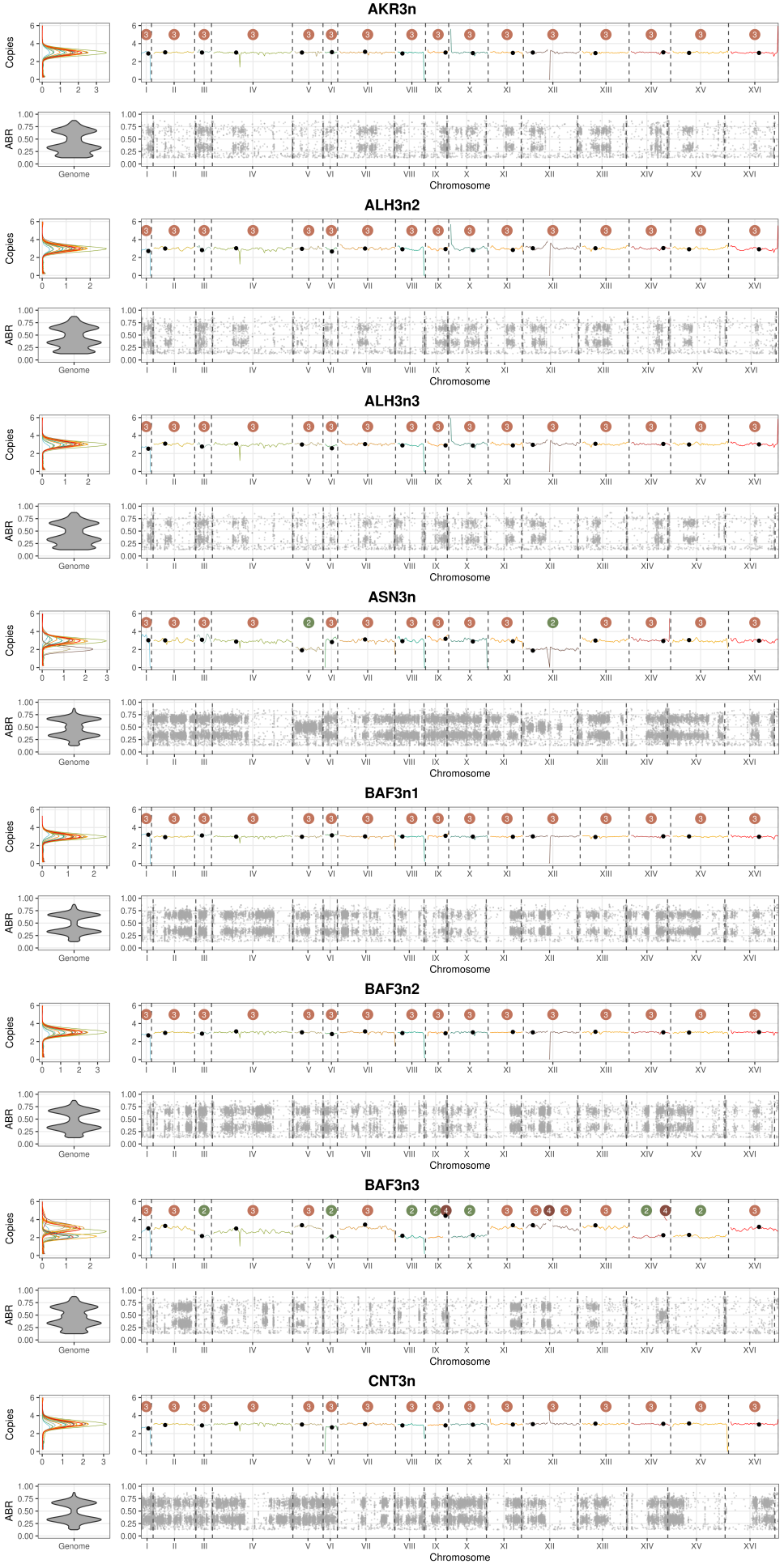

**Figure S6.** Whole-genome profiles of spontaneous triploids recovered following intra-ascus mating of spores derived from diploid strains. Sequencing depth histograms (top left) were used to estimate chromosome copy number (top right; values indicated above each chromosome). Centromeres are marked with black circles. Allele balance ratio (*ABR*) profiles are shown at the genome-wide level (bottom left) and for individual chromosomes (bottom right), with each point representing an alternative allele at a heterozygous site. On three-copy chromosomes, *ABRs* cluster around ~0.33 and ~0.67, corresponding to 1:2 and 2:1 allelic ratios and supporting the triploid chromosome copy number.

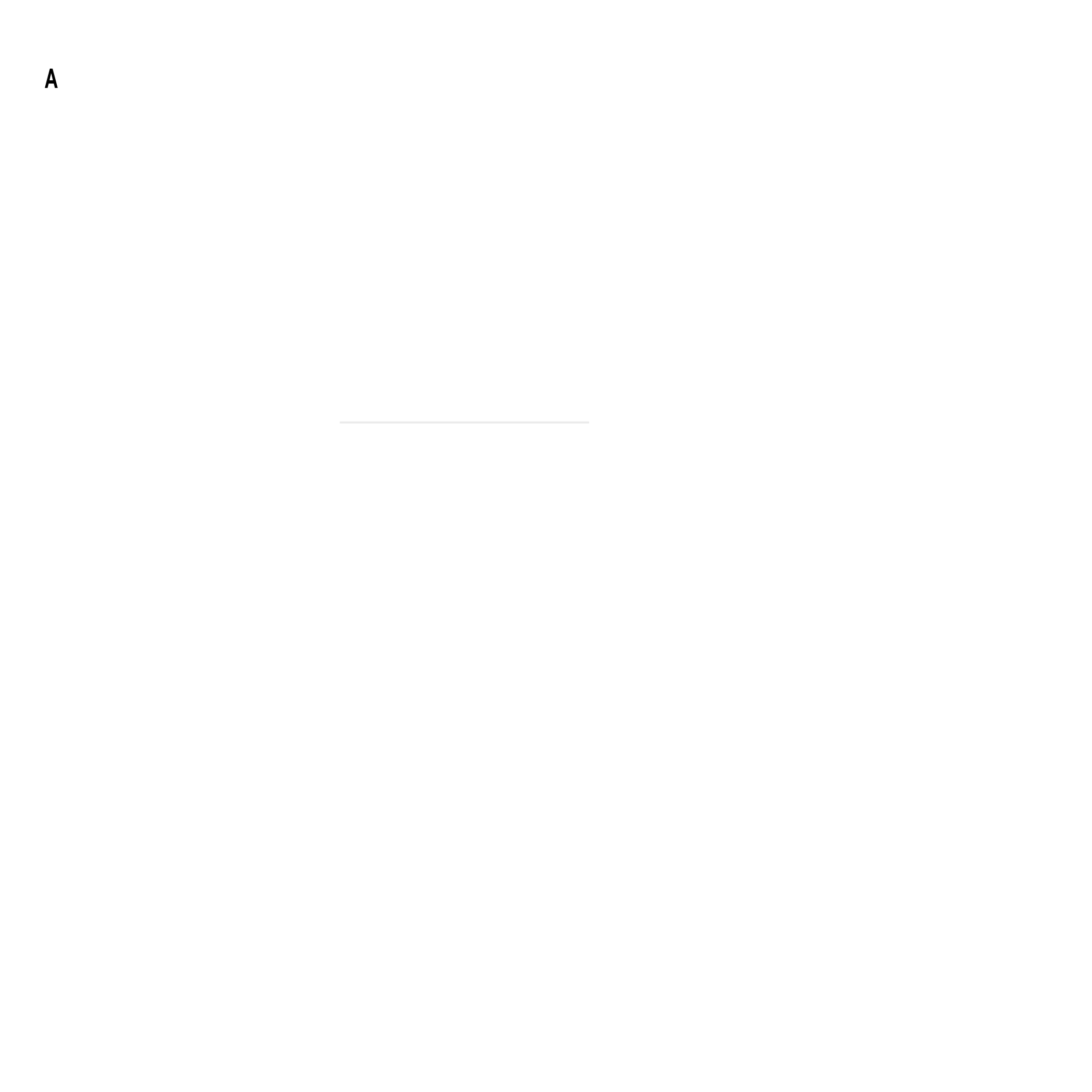

**Figure S7.** A, Inferred ploidy distributions of spore-derived isolates; shaded areas indicate the expected ploidy range of endoreplicated isolates, which was not observed. B, Chromosome copy-number profiles of spore-derived isolates from tetraploid and pentaploid strains. No evidence of post-meiotic endoreplication was detected.

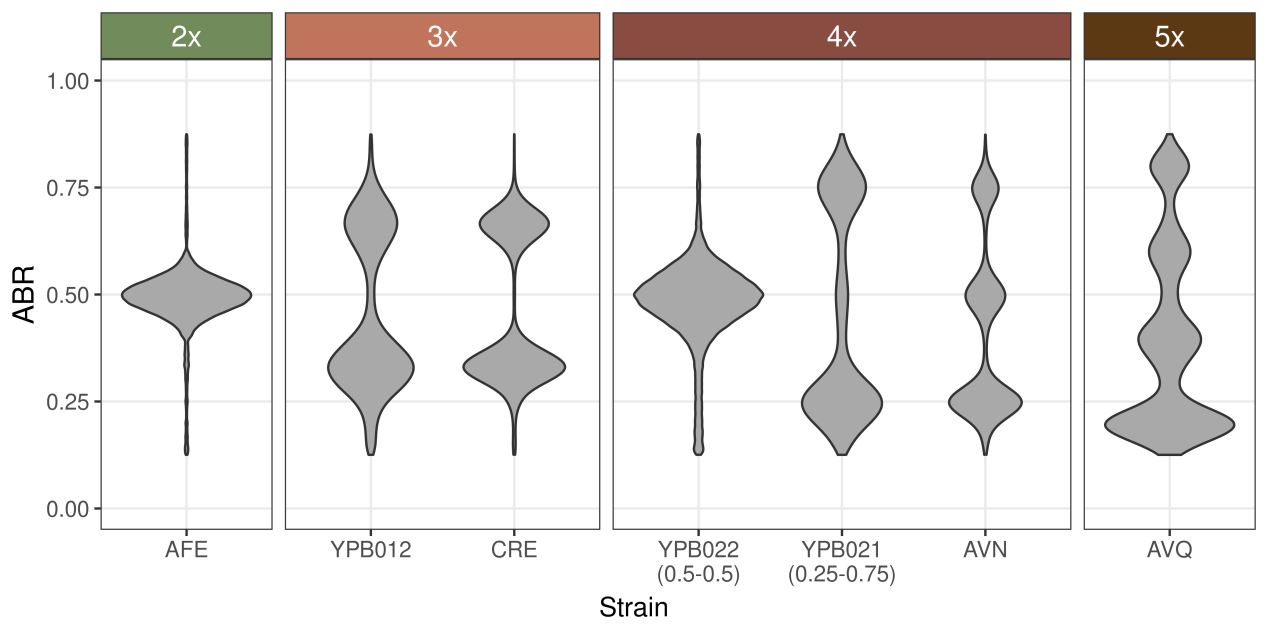

**Figure S8.** Allele balance ratio (*ABR*) profiles of *Saccharomyces cerevisiae* strains with varying ploidy. Only alternative alleles are plotted. Diploid heterozygous strains exhibit a single *ABR* value of 0.5. Polyploids display characteristic discrete *ABR* values: multiples of 0.33 for triploids, 0.25 for tetraploids, and 0.2 for pentaploids. Notably, the *ABR* profiles of natural tetraploids differed from those of artificial tetraploids, presenting all combinations of 0.25 multiples.
