## Supplementary material for "A postmeiotic route to polyploidy": Extented methods

### Extended Methods

#### Ploidy analysis by flow cytometry details

Cellular ploidy was estimated by propidium iodide (PI) staining coupled with flow cytometry. For ploidy validation experiments, strains were grown in 5 mL YPD at 30 °C and processed as previously described(Gómez-Muñoz & Fischer, 2025). For rapid ploidy assessment, either overnight 1 mL YPD microcultures were prepared prior to fixation, or individual colonies were directly fixed in 70% ethanol.

Fixed cells were stained with PI (50 µg mL^-1^) and analyzed using a MACSQuant VYB flow cytometer. Data were processed using CytoExploreR (https://github.com/DillonHammill/CytoExploreR). Initial gating was performed using forward scatter area (FSC-A) versus side scatter area (SSC-A) to exclude debris, followed by FSC-A versus FSC-H to eliminate doublets. To mitigate cell aggregation, two complementary strategies were employed. First, selected samples were subjected to a proteinase K treatment (5 mg mL^-1^) for 2 h at 50 °C prior to staining. Second, an additional gating step using FSC-A versus PI fluorescence intensity (FL4-A; 605-625 nm) was applied. Samples in which more than 50% of events fell within this third gate were considered to yield reliable ploidy estimates. Information regarding proteinase K treatment and gating quality metrics is reported in Table S2 (columns “Proteinase” and “Cells_gate”).

G0/G1 and G2 PI fluorescence intensity peaks were quantified and summarized using MuPETFlow (v0.1.1)(Gómez-Muñoz & Fischer, 2025). Both decimal and rounded ploidy values were extracted and used for downstream analyses.

#### Mass mating assays details

Query and tester strains were grown on YPD agar prior to mating. Using a sterile 1-µL inoculation loop, a small amount of cells from each query and tester strain was streaked separately onto defined 1 cm^2^ areas of YPD agar plates. Approximately half of the cells from each strain were then mixed together in the 1 cm^2^ area between the two streaks, while portions of each culture were left unmixed and served as internal controls.

Plates were incubated for 24h at 30 °C to allow mating. A panel of selective media was used as described in the Media composition and growth conditions section. After incubation, cells from the mixed mating zone were collected using a sterile toothpick and streaked onto double-selection medium (CSM - Ura + G418) using the three-streak dilution method to obtain isolated colonies. In parallel, tester strains were streaked onto single-selection media (CSM - Ura and CSM + G418) to verify marker compatibility (uracil prototrophy and G418 sensitivity) and growth.

Plates were incubated at 30 °C and examined after 3 days. Crosses were scored as positive when multiple colonies were observed on the double-selection medium beyond the initial inoculation streak.

To enable higher-throughput detection of diploid maters, the standard mass mating protocol was modified. Tester and query strains were first resuspended in liquid YPD, mixed, spotted onto YPD agar, and incubated at 30 °C for the same duration as in the standard assay.

After mating incubation, cells were not streaked. Instead, a small amount of cells was collected using a sterile 1-µL inoculation loop, resuspended in 1 mL sterile H_2_O, and diluted 1:10. Diluted suspensions were then spotted onto double-selection medium.

#### Zygote formation and creeping assays

For the zygote formation assay, a small amount of cells from the query strain was mixed with tester strains and incubated for 4 h. Samples were then examined by light microscopy for the presence of zygotes, identified as buds emerging from two fused cells.

Creeping assays were performed as described previously(Arras *et al*, 2022). Briefly, 100 µL of query and tester cell suspensions adjusted to an optical density of 0.2 were mixed in flat-bottom 96-well plates and incubated for 16 h. Creeping behavior was scored based on the formation of a growth ring along the wall of the well.

#### Mating score calculation

Mating efficiency was first assessed semi-quantitatively using the following scoring scheme: no growth (score 0), poor growth (1–10 colonies; score 1), moderate growth (11–50 colonies; score 2), sparse cell layer (score 3), and confluent cell layer (score 4). For each strain, results were recorded as a pair of scores corresponding to crosses with the *a* and *α* tester strains, respectively. Accordingly, a result of (0/4) indicates no mating with the *MATa* tester but robust mating with the *MATα* tester, consistent with an *a* mater phenotype. Semi-quantitative mass mating scores were then converted into partial scores of -1, 0, or 1. Scores below 3 were assigned -1, scores equal to 3 were assigned 0, and only a score of 4 was assigned 1, to restrict the analysis to strains displaying clear mating signals. Zygote formation assays were scored as 1 for positive and -1 for negative outcomes. For creeping assays, positive results were assigned a score of 1, whereas negative results were assigned 0, as weak positive signals may not be reliably detected by this assay. When assays were performed in replicate, partial scores from each replicate were summed and averaged. Discrepant replicates therefore yielded intermediate decimal values. Final mating scores were obtained by summing all partial scores as follows:

| Mass mating | | Zygote formation | | Creeping | |
| --- | --- | --- | --- | --- | --- |
| Result | Partial score | Result | Partial score | Result | Partial score |
| 4 | 1 | Positive | 1 | Positive | 1 |
| 3 | 0 | Negative | -1 | Negative | 0 |
| 2 | -1 |  |  |  |  |
| 1 | -1 |  |  |  |  |
| 0 | -1 |  |  |  |  |

#### Quantitative mating assay

Mating efficiency was quantified using an assay adapted from(de Barros Lopes *et al*, 2002). Query and tester strains were grown on YPD agar, resuspended in liquid medium, and optical density at 600 nm was measured. For each cross, a volume corresponding to 2.5 × 10^8^ cells of each strain was collected by centrifugation and resuspended in 125 µL of sterile 1× PBS.

Equal volumes of query and tester cell suspensions were mixed to obtain a final volume of 250 µL per mating mix. Each mating mix was briefly sonicated (20% amplitude, 15 s) to disrupt cell aggregates. Nitrocellulose filters (7 mm diameter) were placed onto YPD agar, and 5 µL of each mating mix was spotted onto the corresponding filter. Filters were incubated for 4 h at 30 °C to allow mating.

After incubation, each filter was transferred into 1 mL of sterile 1× PBS and vortexed to resuspend cells. Serial dilutions were prepared and plated onto double-selection medium (CSM - Ura + G418) to quantify mating products. Plates were incubated at 30 °C for 3-4 days, after which colonies were counted to calculate colony-forming units (CFUs).
